# **MARGE:** Mutation Analysis for Regulatory Genomic Elements

**DOI:** 10.1101/268839

**Authors:** Verena M. Link, Casey E. Romanoski, Dirk Metzler, Christopher K. Glass

**Affiliations:** Department of Cellular and Molecular Medicine, University of California, San Diego, San Diego, United States; Department of Medicine, University of California, San Diego, San Diego, United States; Division of Evolutionary Biology, Faculty of Biology, Ludwig-Maximilian Universität München, Planegg-Martinsried, Germany; Department of Cellular and Molecular Medicine, University of Arizona, Tucson, United States

## Abstract

Cell-specific patterns of gene expression are determined by combinatorial actions of sequence-specific transcription factors at *cis*-regulatory elements. Studies indicate that relatively simple combinations of lineage-determining transcription factors (LDTFs) play dominant roles in the selection of enhancers that establish cell identities and functions. LDTFs require collaborative interactions with additional transcription factors to mediate enhancer function, but the identities of these factors are often unknown. We have shown that natural genetic variation between individuals has great utility for discovering collaborative transcription factors. Here, we introduce MARGE (**M**utation **A**nalysis of **R**egulatory **G**enomic **E**lements), the first publicly available suite of software tools that integrates genome-wide genetic variation with epigenetic data to identify collaborative transcription factor pairs. MARGE is optimized to work with chromatin accessibility assays (such as ATAC-seq or DNase I hypersensitivity), as well as transcription factor binding data collected by ChIP-seq. Herein, we provide investigators with rationale for each step in the MARGE pipeline and key differences for analysis of datasets with different experimental designs. We demonstrate the utility of MARGE using mouse peritoneal macrophages, liver cells, and human lymphoblastoid cells. MARGE provides a powerful tool to identify combinations of cell type-specific transcription factors while simultaneously interpreting functional effects of non-coding genetic variation.

## Introduction

Molecular mechanisms enabling cell-specific transcriptional responses to intra- and extra-cellular signals remain poorly understood. Genome-wide studies of most lineage-determining (LDTF) and signal-dependent transcription factors (SDTF) indicate that the vast majority of their binding sites are in distal intra- and intergenic locations that frequently exhibit epigenomic features associated with enhancers (1–6) and are evolutionary well conserved (7–9). The complement of active *cis*-regulatory elements bound by LDTFs changes across cell types, whereas promoters stay the same. Therefore, these findings introduced the notion that enhancers are largely responsible for cell type-specific gene expression (10–12). The ENCODE consortium annotated epigenetic features associated with enhancers in several different cell lines, primary cells and tissues providing evidence for hundreds of thousands of such elements in the human genome (13), greatly exceeding the number of promoters.

Previous studies of macrophages and B cells provided the basis for a collaborative and hierarchical model (14–16). In this model, collaborative binding of two or more LDTFs opens up chromatin to establish enhancers (1), enabling cell-specific actions of broadly expressed SDTFs (17) (reviewed in (18)). The collaborative nature of LDTFs was further demonstrated by analysis of effects of genetic variation in macrophages provided by two inbred strains of mice (19).

Genome-wide association studies, or GWAS (20) have shown that most complex trait-associated genetic variation is located in non-gene/protein regions of the genome. Such noncoding variants have the potential to change conserved sequences recognized by LDTFs and thereby alter enhancer landscapes between different alleles. These differences could manifest between individuals (i.e., between individuals that are each homozygous for opposite alleles), or within an individual that is heterozygous for a functional enhancer variant. A straightforward mechanism by which enhancer function would be altered by genetic variation is where alleles alter the affinity of transcription factors to bind their motifs. Consistent with the enhancer model whereby transcription factors collaborate with each other to bind DNA motifs, reports have found that allelic variation that mutates DNA binding motifs reduces binding of the respective factor while at the same time reducing binding of collaborating factors within 100 base pairs (19,21,22). Since the DNA binding motif of the partner factor is not mutated, these examples demonstrate a coordinated action of transcription factors in accessing DNA. The implication for cell-specific gene regulation is that genetic variants altering collaborative factor binding at enhancers will only be functional in the appropriate cell type where the correct combinations of transcription factors are expressed. The practical implication of these observations is that the particular combinations of factors may be discovered with the general strategy in any cell type. In addition to the discovery of transcription factors, this method identifies the precise genomic loci where genetic variation has a functional role in factor binding that may influence higher order biological processes.

To facilitate discovery of novel collaborating transcription factors using the genetic variation approach, we developed MARGE (**<u>M</u>**utation **<u>A</u>**nalysis for **<u>R</u>**egulatory **<u>G</u>**enomic **<u>E</u>**lements). MARGE is a suite of software tools to analyze ChIP-seq, ATAC-seq, DNase I Hypersensitivity or other next generation sequencing (NGS) assays where genotyping or DNA sequence data is available.

MARGE requires two data types: 1) genetic variation, and 2) high-throughput sequencing data (ChIP-seq, ATAC-seq, DNasel-seq). MARGE then integrates these data and provides visualization tools to interpret the results. Importantly, MARGE was built to test for functional effects of alternate alleles at single nucleotide polymorphisms (SNPs) as well as short insertion-deletions (InDels). MARGE performs traditional de-novo motif analysis on genomic sequence for each polymorphic allele to identify DNA binding motifs that potentially affect transcription factor binding based on sequence analysis alone. The next step is to test whether the set of potential variants that mutate a single DNA binding motif are enriched in a set of loci where differential binding/accessibility is observed. For this step, MARGE associates quantitative measures of binding or accessibility from the ChIP/ATAC/DNaseI-seq data with the list of potential mutations in motifs. It analyzes differences in two genotypes by comparing the transcription factor binding distribution in relation to motif mutations between both genotypes, and also takes advantages of a Linear Mixed Model (LMM) (23,24) to compare many different individuals at the same time

In this report, we apply MARGE and demonstrate its ability to reliably identify known key regulators of macrophage lineage. We further apply MARGE to three different ChIP-seq datasets from mouse liver cells and also show that MARGE can identify important B cell factors in a human PU.1 ChIP-seq dataset from lymphoblastoid cell lines (25). In conclusion, MARGE is the first publicly available tool that is created to identify combinations of collaborating transcription factors. This approach is agnostic to cell type and can be applied in any dataset where genotypes and epigenetic signatures are measured.

MARGE is based on the ChIP-seq analysis tool HOMER (1) (http://homer.ucsd.edu/homer/) and it is an extension to the software used in (19,21). The source code and installation package are freely available on GitHub: https://github.com/vlink/marge/blob/master/MARGE_v1.0.tar.gz

## MATERIAL AND METHODS

### Overview

A schematic outlining the major steps of MARGE is shown in Fig 1. First, MARGE offers a complete pipeline to process VCF (Variant Call Format) files (26) and generate individualized diploid genomes by extrapolating genetic variants from VCF files and swapping in alternate alleles into a reference genome (Fig. 1a-b). Importantly, MARGE is able to analyze sequencing data from homozygous (e.g., inbred mouse strains) and heterozygous (e.g., human) genomes and includes analysis for Single Nucleotide Polymorphisms (SNPs) as well as short Insertion-Deletions (InDels). Because MARGE generates genomes for each individual in the VCF file, the investigator can map their sequencing data to the genome with all genetic variations by using user-defined mapping software (e.g. bowtie2 (27) or STAR (28)) (Fig. 1c). MARGE shifts positions of individual sequence to their corresponding reference coordinates for motif analysis and visualization (Fig. 1 d-g). MARGE offers de-novo motif analysis for individualized genomes (Fig. 1g), as well as a new algorithm to identify transcription factor binding motifs associated with allele-specific transcription factor binding or open chromatin (Fig. 1h). Each step of the MARGE pipeline is discussed below.

**Figure 1:**
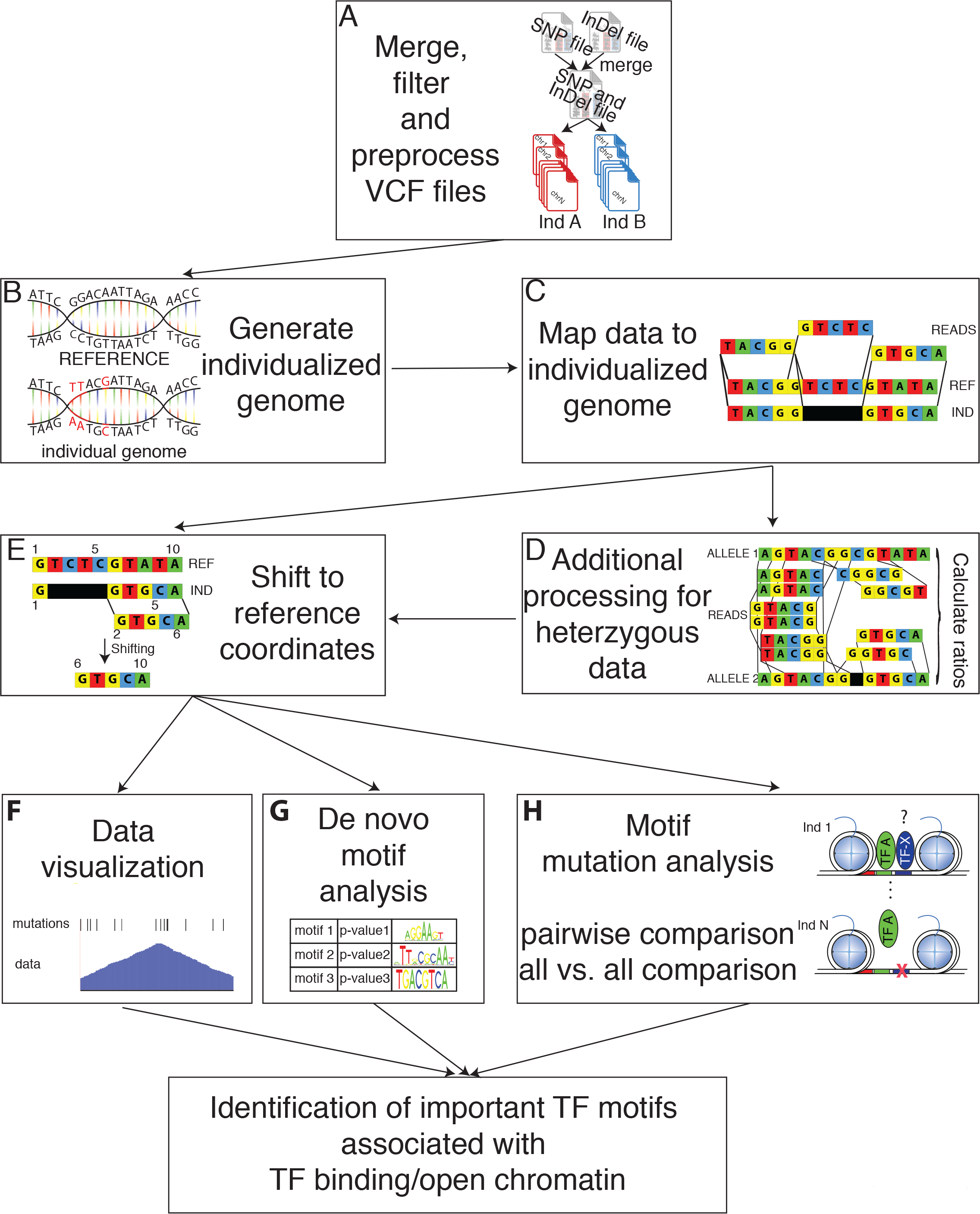
Overview of the MARGE pipeline. (**A**) MARGE merges VCF files for SNPs and InDels, offers some basic filtering and split the merged VCF file into separate genotype-specific mutation files. (**B**) It then generates individual genomes by inserting the annotated mutations in the reference genome per genotype and (**C**) allows mapping of the experimental data sets to the individualized genomes. (**D**) The data mapped to the individualized genomes is then shifted back to the reference coordinates. (**E**) In case of heterozygous data additional processing is necessary. MARGE offers (**F**) scripts for data visualization including BED files for genetic variation per genotype. It further offers (**G**) de-novo motif analysis for the individual genomes to make sure the enrichment analysis is performed on the correct sequence instead of the reference. MARGE also offers a new algorithm (**H**) to associated TF binding motifs with genotype-specific binding for pairwise comparisons, as well as comparisons for many different individuals (all-versus-all comparison). Taken all of that together MARGE is able to identify TF binding motifs that are functionally associated with TF binding.

### Merge, filter, and pre-process VCF files

The initial step of the MARGE pipeline is to generate a set of high-confidence sequence differences between the alleles of interest (Fig. 1a). MARGE allows some basic filtering of VCF files by quality scores, however VCFtools (26) provides more sophisticated tools for this purpose. For some sequencing projects like the mouse genome project (29), SNPs and InDels are annotated in separate files, whereas other projects like the 1000 Genome project (30) provides one large file with SNPs and InDels. When SNPs and InDels are provided separately, MARGE merges them as a first step. If a combined file is provided then the first processing step is skipped. In cases where SNPs overlap deletions or insertions within one genomic background the SNP is filtered out and the longer mutation is kept. MARGE also simplifies the annotation of the variants per genotype (Fig. 2a). In cases where more than one possible mutation occurs in a particular genomic location (e.g. two different genotypes have two different mutations in comparison to the reference genome), the mutation is not always annotated as the shortest mutation per genotype. As shown in Fig. 2a the genetic variant for genotype2 is annotated as GTT -> GTTGTT. MARGE processes each genotype separately and therefore calculates the shortest genetic variation for each genotype (in this case T -> TGTT).

### Generating individualized genomes

MARGE produces individualized genomes by inserting the alleles from the VCF file into the reference genome and generating fasta files, which then can be used to make indices for mapping software. For homozygous data, only one genomic sequence is generated. Generation of individualized genomes and interpretation of allele-specific mapping for heterozygous data requires an additional step. Specifically, alleles at heterozygous sites need to be assigned on the same chromosome as neighboring heterozygous alleles. In genetics, this is called knowing the *phase* of the genotypes. Phase is especially important for MARGE when variants are in close proximity, because most sequencing reads are between 50-200 base pairs in length. When multiple SNPs reside in the same read, the correct combination of alleles in the genomic index is essential for accurate mapping and downstream interpretation. MARGE inherently assumes that all heterozygous data is phased. There are good resources for phasing genotypes in human populations. For example, phasing can be achieved using BEAGLE (31) or SHAPEIT (32) in conjunction with known haplotype structure of large reference populations such as the 1000 Genomes Project. In cases were phasing is not easily possible (e.g. F2 generation of inbred mice) loci where mutations overlap within the read length should be excluded from the analysis.

### Mapping data to individualized genomes

Mapping of sequencing experiments to the individualized genome provides better results and decreases the possibility of incorrect mapping due to technical bias (Fig. 1c). This is especially true in datasets with a large number of differences to the reference. In these cases, mapping to the reference can introduce bias and in the case of datasets containing heterozygous genotypes can lead to overestimation of allele-specific expression or binding (33–35). To assess the effect of individualized genomes on mapping, we used a ChIP-seq dataset from inbred strains of mice. This provided a simplified situation since their genomes are entirely homozygous and all sequence tags originated from a genome of known sequence. Specifically, we used a PU.1 ChIP-seq dataset from three strains of mice (C57BL/6J, NOD/ShiLtJ, and SPRET/EiJ) (21). C57BL/6J (C57) is the commonly used reference genome and differs to NOD/ShiLtJ (NOD) in about 5 million genetic variants (89% SNPs, 11% InDels), whereas SPRET/EiJ (SPRET) provides about 43 million variants (89% SNPs, 11% InDels). Mapping of the ChIP-seq data to their respective genomes affected the overall mappability of the reads (Fig 2b) and the percentage of uniquely mapped reads (Fig 2c). The difference in mapping is directly correlated to the number of differences between the genomes. After removing all reads that map to multiple locations, peaks were called on all datasets separately and compared. Peaks from the C57 ChIP-Seq mapped to C57 and NOD genomes show only small differences (Fig. 2d) (about 1% of peaks are unique to either genotype), but increasing the number of variation between the genotypes lead to many peaks uniquely called in one of the mapped datasets (up to 12%). Also when comparing a PU.1 ChIP-seq dataset in human lymphoblastoid cell lines (25) mapped to the reference versus the individual genomes only about 90% of reads where mapped to the same loci (Supp. Fig. 1a). The number of differences between the hg19 reference genome and the individualized genomes is smaller than for the mouse data, but still up to 4% of peaks were uniquely called on either the dataset mapped to the reference or the individual genome (Supp. Fig. 1b). Therefore, mapping the data to the correct individualized genome increases the mapping accuracy substantially, leading to a more precise downstream analysis.

**Figure 2:**
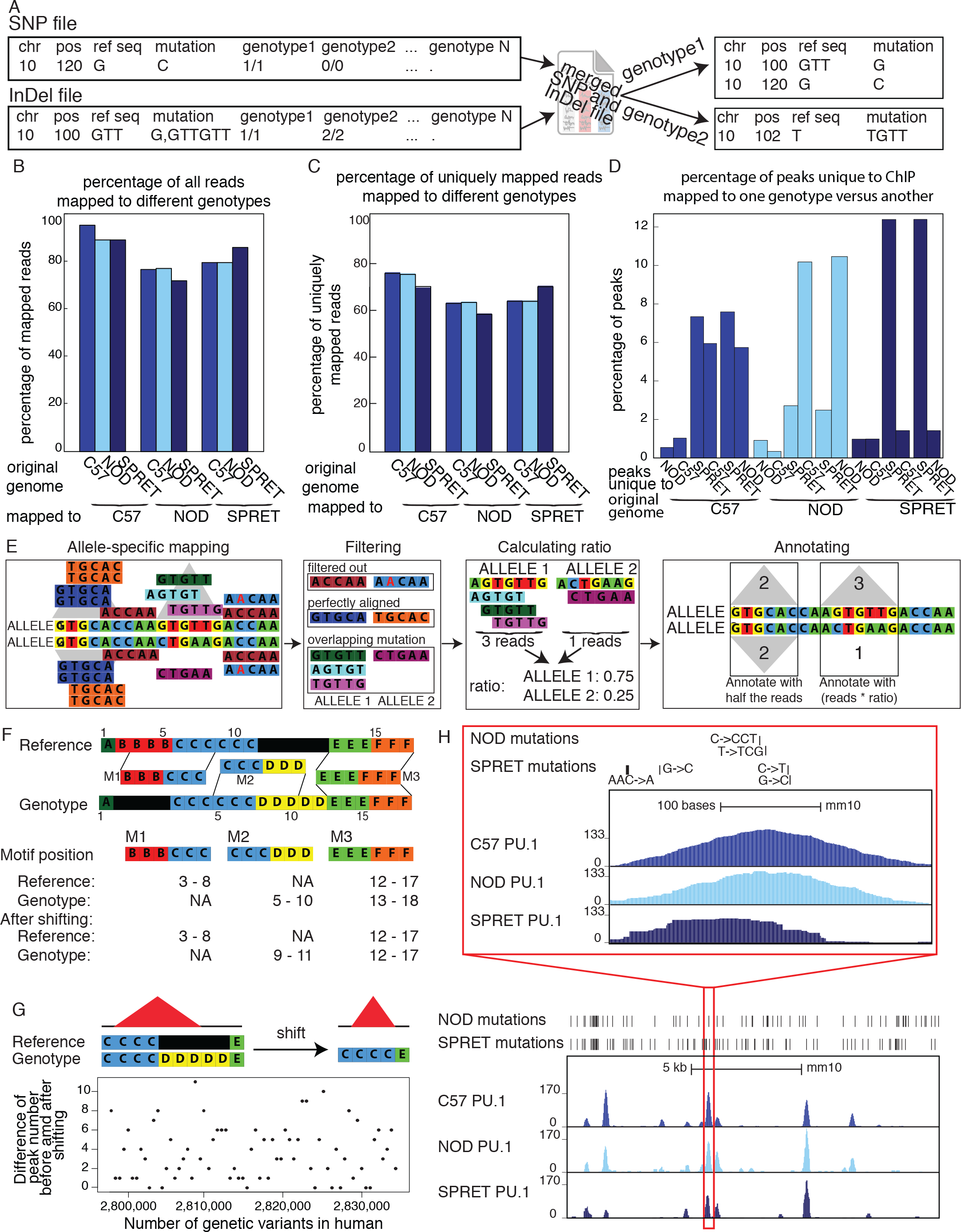
Details of pipeline: (**A**) MARGE merges SNP and InDel VCF files and then splits the merged file. It finds the shortest annotation for each mutation, changing the original annotation from the VCF file. (**B**) Comparison of the overall mapping efficiency. There is a small decrease in overall mappability when data is mapped to the reference. (**C**) Comparison of mapping efficiency for uniquely mapped reads after mapping to different genomes. There is an increase in mapping performance when mapped to individualized genomes. (**D**) Percentage of peaks uniquely called to dataset mapped to one genotype versus another. Up to 12% of peaks are unique to one genotype. (**E**) Pipeline for processing heterozygous data: Data is mapped to both alleles and shifted back to the reference coordinates. Reads that do not uniquely align to the genome are filtered out. Perfectly aligned reads, as well as perfectly aligned reads overlapping mutations are filtered out and peaks are called on perfectly aligned reads. For each locus without any mutations, the peaks for both alleles are annotated with half the reads that mapped to this locus. For each locus with mutations a ratio is calculated based on the reads overlapping mutations and then the locus is annotated with the number of perfectly aligned reads multiplied by the corresponding to the ratio. (**F**) Schematic of the shifting process: Genomic coordinates of the individual genomes do not concur with the reference due to InDels. MARGE shifts the individual coordinates to the reference without changing the length of the sequence. (**G**) Shifting peak coordinates leads to minor loss of peaks. 34 PU.1 ChIP-seq data sets were mapped and peaks were called before and after shifting. Even with 2 million genetic variants between the reference and the individualized genomes only up to 11 peaks are different. (**H**) UCSC genome browser shot showing PU.1 ChIP-seq data in large peritoneal macrophages in 3 different inbred strains of mice (C57, NOD, and SPRET). Bed graphs generated by MARGE show genetic differences between the strains. The red rectangle shows a zoomed-in area of the UCSC genome browser.

### Additional processing for heterozygous data

Many studies in mice use hybrid mouse strains (F1) generating heterozygous mice from two homozygous parents (Fig 1d). Furthermore, all human genomes are heterozygous in many loci and due to the advantages in sequencing technology, have become more realistic to study genome-wide. To improve mapping for heterozygous data, statistical methods have been developed (e.g. WASP (36)). Unfortunately, these methods can only handle SNPs. In order to also analyze heterozygous data with InDels, we map our data to two reference genomes corresponding to alternative parental alleles. To effectively analyze heterozygous data, allele-specific expression or binding needs to be calculated. For this step, MARGE filters all reads with perfect alignment followed by filtering of all reads spanning a variant between the two parental strains (Fig 2e). If the heterozygous data is not phased, all regions that contain more than one mutation within the length of one read should be excluded from the analysis. This procedure makes sure that it is possible to confidently identify the allele of origin. To assign allele-specific reads correctly all loci without any variation are annotated with half of the perfectly aligned reads, because half of the reads that are sequenced originate from allele 1 and the other half from allele 2. For loci with allele-specific sequences, the ratio of reads per allele is calculated based on the reads spanning variations. Then the loci are annotated with the corresponding ratio of all perfectly aligned reads mapped to this locus.

### Shifting to reference coordinates

A major challenge of mapping data to individual genomes is that the experiments cannot be easily compared because of insertions and deletions (Fig. 1e). For example, the chromosomal locations between individuals (and across homologous chromosomes within heterozygous individuals) do not correspond to each other anymore. Therefore, to be able to use external analysis software and to visualize the data in the UCSC genome browser (37), we designed MARGE to shift mapped data back to reference coordinates (Fig. 2f). To accomplish this, MARGE generates shifting vectors for each genome (or haploid genome in the case of human/heterozygous data). Motifs can overlap insertions (M2) and deletions (M1) in the reference genome (Fig. 2f). The M2 motif consists of 6 bases, but after shifting the length is shrank to 3 bases due to the deletion. Therefore, positional shifting has the potent to introduce problems. For example, InDels can cause potential TF binding motifs to disappear or appear, which is of interest because these cases likely have functional consequence. Another complication of shifting coordinates occurs in the identification of ChIP-seq peaks from variable chromosomal sequences (i.e. shifting can cause a loss of peaks). This is because ChIP-seq peak calling tools often require a minimum length in order to identify peaks and this might not be reached after shifting. To check how frequently a peak was lost, each PU.1 ChIP-Seq dataset performed in human lymphoblastoid cell lines (25) was mapped to its individual genome and peaks were called with HOMER both before and after shifting. There are up to 2 million genetic differences between the reference genome (hg19) and the allele-specific genomes per individuals, but only up to 11 peaks are lost after shifting (which corresponds to less than 0.1% of all peaks) (Fig. 2g, Sup. Table 1). Also when repeating this procedure for diverse mouse strains (with more than 40 million genetic differences) only about 0.2% of all peaks were lost (Sup. Table 2). Therefore, despite an opportunity for difference to emerge in peak calling, we conclude that this phenomenon is very rare and does not offset the advantages from more accurate mapping.

### Data Visualization

Tools like the Integrative Genomics Viewer (IGV) (38,39) allow visualization of individual genomes, but require the user to install the software locally, which is not preferable for data sharing. One of the most common software platforms to visualize next-generation sequencing data online is the UCSC genome browser (37). Although a powerful tool, it does not allow the usage of other genomes than the references. To account for this, after shifting the genomic coordinates from the individualized genomes to the reference genome, MARGE can generate UCSC genome browser files (e.g. bedGraphs and bigWig files) that take into account individual genomic features (Fig. 1f). In addition, MARGE can generate BED (Browser Extensible Data) (40) files with all alternate alleles relative to the reference coordinates for upload to the genome browser (Fig. 2h). We also provide basic tools to interact with the different individual genomes. For example, we make it possible to directly compare the number of polymorphisms between different datasets in a table format for either all variants (Table 1) or for all private variants (those which can only be found in a particular individual compared to all others) (Table 2). More importantly however, MARGE can align nucleotide sequences from different individuals or chromosome sequences such as nucleotides or protein sequences. This application integrates RefSeq (41) or common gene name information to provide alignment for genes of interest, but is also able to extract the sequence for every genomic location of interest. This provides a fast and easy way to check for differences in genes or non-coding regions for different genetic backgrounds. This also simplifies the design of primers or other constructs, because differences can be checked by simple alignments of VCF files. To enable some more user-specific analysis, MARGE annotates files containing genomic coordinates with all genetic variants and generates files with genotype-specific sequences.

**Table 1:**
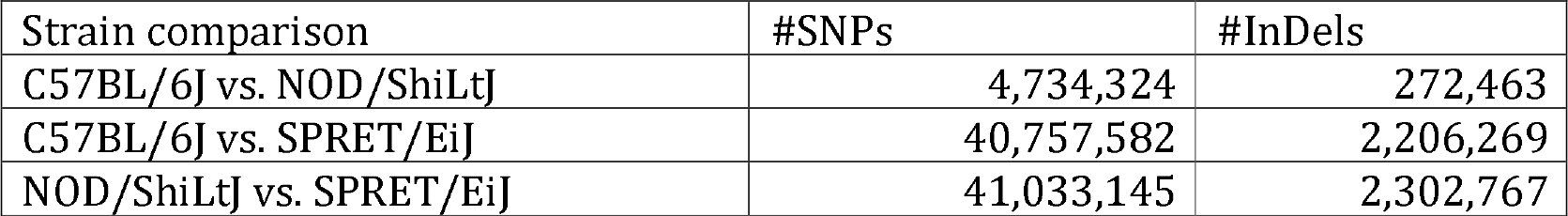
Overview of all natural genetic variation found in all strain-wise comparisons

**Table 2:**
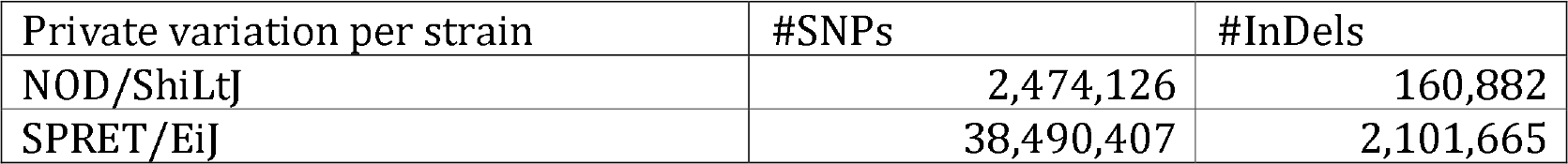
Overview of all private genetic variation found in in this strain versus all other strains

### De novo motif analysis

One of the first steps in analyzing ChIP-seq data is motif analysis. The de-novo motif analysis software from HOMER (1) was adapted to allow the integration of the individual genomes (Fig. 1g). We extended the de-novo motif finding algorithm (1) with a function to extract the sequences of the different genotypes as inputs to make sure that the motif finding algorithm is applied to the correct sequences and finds the motifs enriched in the sequence of the genotype not of the reference. It is possible to use different genotypes for the foreground sequences and the background sequences when unique peaks in two different genotypes are compared as foreground and background. These extensions make MARGE a powerful tool in comparing enriched motifs in two different genotypes.

### Motif mutation analysis

MARGE was primarily developed to determine importance of various nearby transcription factor motifs on the binding of a given transcription factor (Fig. 1h). It can analyze transcription factor binding profiles for two genomes in a pairwise fashion, but is also able to analyze the binding profiles of many different genomes together (Fig. 3). The first case is preferable when two datasets have many genetic differences (e.g. two diverse mouse strains), as it may be more cost effective experimentally (pairwise comparison). For the analysis of human samples, however, it may be preferable to have more individuals, as the number of differences between two human genomes is fewer. In this scenario, a larger sample size may be required to achieve statistical power (all-versus-all comparison). MARGE uses a list of hand-curated motifs from the JASPAR motif database (42) as default, but also allows user-defined input.

**Figure 3:**
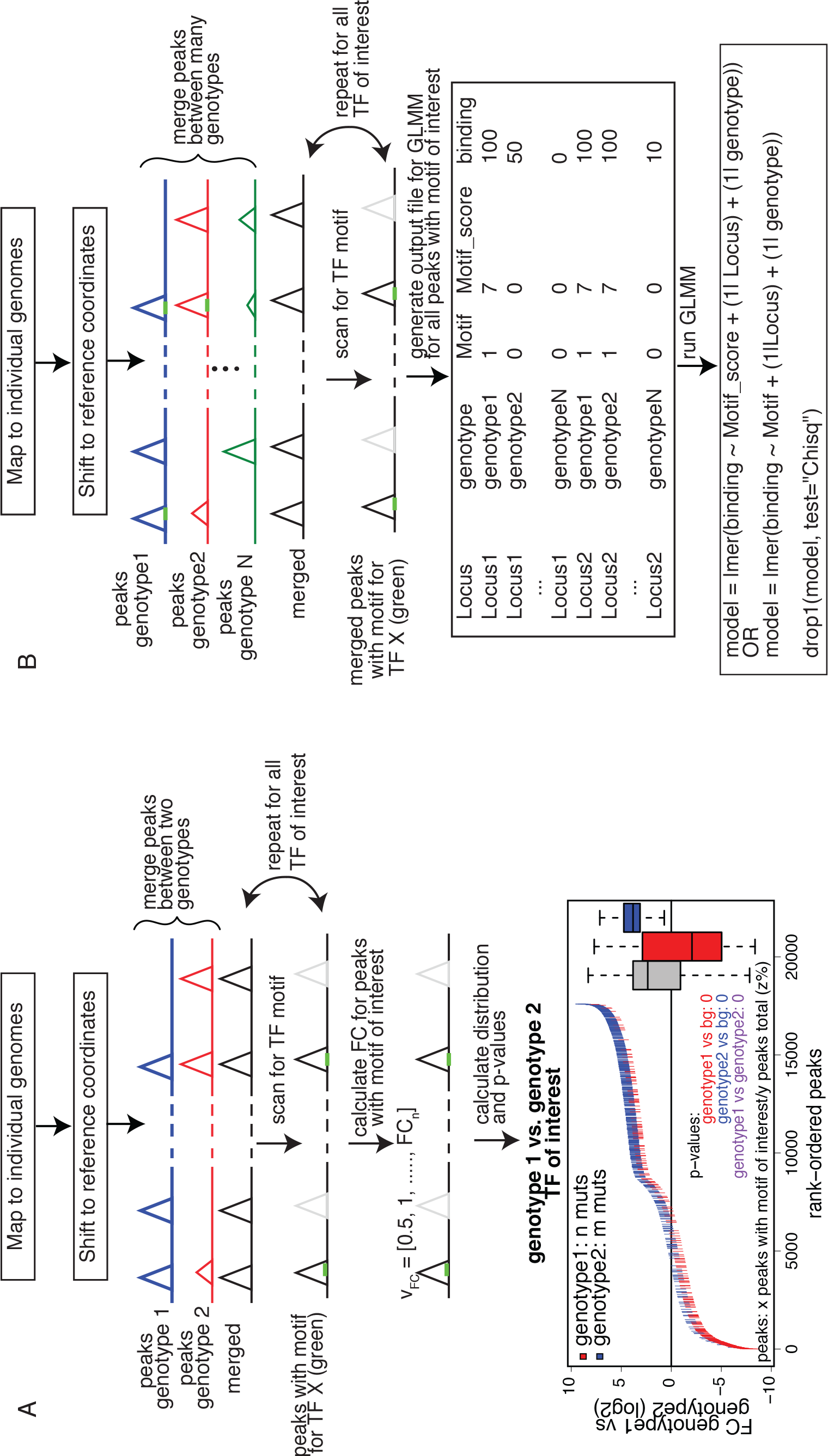
Schematic showing the algorithm for the motif mutation analysis for pairwise comparisons or comparisons of a big group of individuals. (**A**) Pairwise comparison: Data is mapped to individual genomes and shifted to reference coordinates. Peaks are called per genotype and are subsequentially merged and annotated with the tag counts from the tag directories with HOMER. The merged file is iteratively scanned for the TF binding motifs of interest. For all peaks containing the current TF motif of interest (marked in green) the binding difference between the two genotypes is calculated (fold change). For each TF the fold change distribution of all peaks is plotted (more information Sup. Fig. 1a) and a Student’s t-test is performed on the fold change distribution of all peaks versus all peaks containing a mutation in genotype1 (red) (genotype2 (blue), respectively). Further a t-test is performed comparing the fold change distribution of all peaks missing the motif of interest in genotype1 versus genotype2 (purple) and corrected for multiple testing. (**B**) Motif mutation analysis on more than two genotypes: Data is mapped to the individual genomes, shifted back to the reference coordinates and peaks are called on each genotype separately and subsequentially merged and annotated. Heterozygous data should be annotated with MARGE’s annotation function. The merged file is iteratively scanned for the TF motif of interest (marked in green). Per TF an output file is generated containing the locus, the binary existence of a motif, the motif score and the read counts. This output file is then inserted into a linear mixed model (LMM) implemented in R with the package lme4 modeling the binding as dependency of the motif score (or motif existence) with random factors Strain and Locus. A p-value is generated using the R command drop1 and corrected for multiple testing.

#### Pairwise comparison

For the pairwise comparisons, peak files of both genotype alignments are merged and annotated with read counts (Fig. 3a). To account for differences between the alleles, the individual genome sequence is extracted and scanned with the motif-scanning algorithm provided by HOMER. Each motif is analyzed separately. Peaks without the motif that is currently scanned for are excluded from the analysis of this particular motif, but are considered for other motifs. Therefore, the analysis of every transcription factor motif is done on a different number of peaks. The fold change of the normalized read counts between the two alleles is calculated. Finally, the distribution of the fold change is calculated for all peaks, all peaks with a mutation in the motif of interest in allele1 and all peaks with a mutation in the motif of interest in allele2. To ensure that a motif is not just considered allele-specific because its log-odd score was slightly below the arbitrarily defined threshold in one of the alleles, MARGE extracts the sequence of the potential motif from each allele and calculates the log odd score based on the provided position weight matrix (PWM). By default a motif is considered missing when the log odd score is smaller or equal to zero, but the user can change this value to whatever seems suitable. MARGE also provides the possibility to define a motif as missing when its log-odd score in one allele is less than n% of the log-odd score in the other allele. To determine the significance of every motif a Student’s t-test is performed between the general fold change distribution and the fold change distribution of allele1 and allele2, respectively.

Furthermore, the p-value between the distributions of the two alleles is calculated. This procedure is repeated for all transcription factors of interest. All p-values are multiplied by the number of comparisons to correct for multiple testing.

Allele-specific binding can be observed due to the loss of the binding site for the collaborative factors or the measured transcription factor itself. Additionally to analyzing every peak with the motif of interest, MARGE can analyze only peaks where all loci with differences in the motif of the measured TF between genotypes are filtered out. A Student’s t-test is performed on the remaining distributions and the p-values are multiplied by the number of comparisons. MARGE outputs a motif mutation plot showing the distribution of mutations in relation to the fold change for each transcription factor (bottom Fig. 3a, Sup. Fig. 2a). It further outputs a density distribution plot for the fold change distribution of all peaks with changes in the motif in allele1, allele2, and the background (Sup Fig. 2b).

#### All-versus-all comparison

In order to perform an all-versus-all comparison on more than two genotypes, peaks are called for all genotypes individually (Fig. 3b) and annotated with read counts. In case of heterozygous genotypes, peaks should be called on alleles separately and also be annotated with allele-specific reads (Fig. 2e). Both alleles are then analyzed as if they were independent genotypes. Therefore, when comparing for example 3 heterozygous genotypes, MARGE actually analyzes 6 independent samples. All sequences of all genotypes are scanned for the motifs of interest. To model the impact of the motif on the binding of the measured factor a Linear Mixed Model (LMM) is used. The binding of the measured factor is modeled as the fixed effect motif existence or motif score (defined by the user) with random effects locus and genotype (Formula 1) with the lme4 package (43) in R (44).

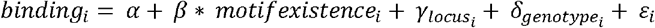

or

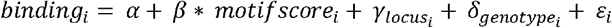

with

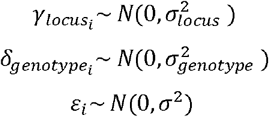

To calculate significance for each motif, the drop1 command is used. It compares a model including motif score (motif existence, respectively) with a model without motif score (motif existence, respectively) and reports the Akaike information criterion (AIC) (45) for the difference. To keep the run time reasonable, MARGE implements threading for this procedure.

### Data mapping

All data was mapped using bowtie2 (27) with default parameters. The data for the different inbred strain of mice and the human data were mapped to the individualized genomes. The individualized genomes were generated using bowtie2-build with default parameters. The data for C57BL/6J was mapped to the mm10 reference genome from the UCSC genome browser (37). The human reference genome was hg19. Uniquely mapped reads are all reads that were mapped to only one unique region of the genome.

To analyze the impact of the genome on the accuracy of the mapping, all mouse ChIP-seq data sets in LPMs (21) were mapped to the three strain genomes C57, NOD, and SPRET. For the human data (25) all data was mapped against the individualized genome for allele 1, allele 2 and the hg19 reference genome. To assess the impact of the mapping on peak calling all reads that were mapped to more than one region of the genome were removed.

### ChIP-seq analysis

All ChIP-seq data sets were analyzed with HOMER after being shifted to reference coordinates. Peaks were called using findPeaks with default parameters and -style factor. For the LPM data set inputs were used for the peak calling. In case of the liver data and the human data no input was available and peaks were called without inputs. After running MARGE on the data, the list of significant motifs was reduced and summarized using HOMER’s compareMotifs.pl.

### Simulation of a data set

MARGE is based on the model of collaborative binding for TFs and important collaborative TF binding motifs therefore should be identified as significant. According to this model a TF can only bind if the collaborative factor can bind, too. Applying this idea to two different genotypes means that if the motif is missing in genotype1 the binding of the measured factor should be lost in genotype1 and be not affected in genotype2 (genotype-specific binding). It further means if the motif is found in both genotypes binding should be similar between them (genotype-similar binding).

For the synthetic dataset, ten motifs were randomly chosen and defined as important collaborative TF for PU.1 (Tead3, Ventx, and Zic1), somewhat collaborative (Rora, Znf354c, and Plag1) and not collaborative (Pax6, Nr4a2, Lin54, and Bhlha15) (Fig. 4a). The genomes from three mouse strains (C57BL/6J (C57), BALB/cJ (BALB), and SPRET/EiJ (SPRET)) were scanned for the occurrence of all motifs (including PU.1). Next a peak file was generated for all genomic locations where the motif of interest was within 200bp of the PU.1 motif. These files were merged between two strains (C57 and BALB, C57 and SPRET, BALB and SPRET). To model genotype-specific binding, the fold change was randomly chosen to be between 2 and 10fold. For genotype-similar binding the fold change between the strains was within 1.5 fold. In all cases the read counts were randomly chosen between 0 and 500. To include biological noise in this dataset 85% of peaks with genotype-specific TF binding motifs follow the genotype-specific binding for highly collaborative motifs. For somewhat collaborative motifs 50% follow this pattern, whereas in the case of not collaborative motifs only 10% of peaks with genotype-specific TF motifs also show genotype-specific binding. To model genotype-similar binding for all highly collaborative motifs 85% of all peaks with the same motif show genotype-similar binding, for somewhat collaborative motifs 50% of the peaks have genotype-similar binding, whereas for not collaborative motifs only 10% show genotype-similar binding. The rest of the peaks show genotype-specific binding randomly assigned to one of the two strains.

## RESULTS

### MARGE recognizes collaborative motifs in synthetic dataset

To test the accuracy of the method, a synthetic dataset was generated simulating a ChIP-seq experiment using an antibody against PU.1 (for more details see Material and Methods, Fig. 4a). Ten motifs were randomly chosen and defined as important collaborative TF for PU.1 (Tead3, Ventx, and Zic1), somewhat collaborative (Rora, Znf354c, and Plag1) and not collaborative (Pax6, Nr4a2, Lin54, and Bhlha15) (Fig. 4a). Data was simulated for three different homozygous mouse strains (C57, BALB, and SPRET). Comparing one representative of the different motif categories shows that the algorithm is able to detect very high significance for Tead3 (defined as highly collaborative), medium significant for Plag1 (defined as somewhat collaborative) and no significance for Nr4a2 (defined as not collaborative) (Fig 4b, Sup. Fig. 2a). In all three comparisons about 50% of all peaks had the motif of interest, so the significance is not dependent on the percentage of peaks having the motif. The algorithm is able to detect significance for all motifs that were collaborative and showed lower or no significance for all non-collaborative motifs (Fig. 4c). PU.1 was almost always recognized as a significant motif, which is expected as the peaks were modeled according to a PU.1 ChIP-seq experiment.

**Figure 4:**
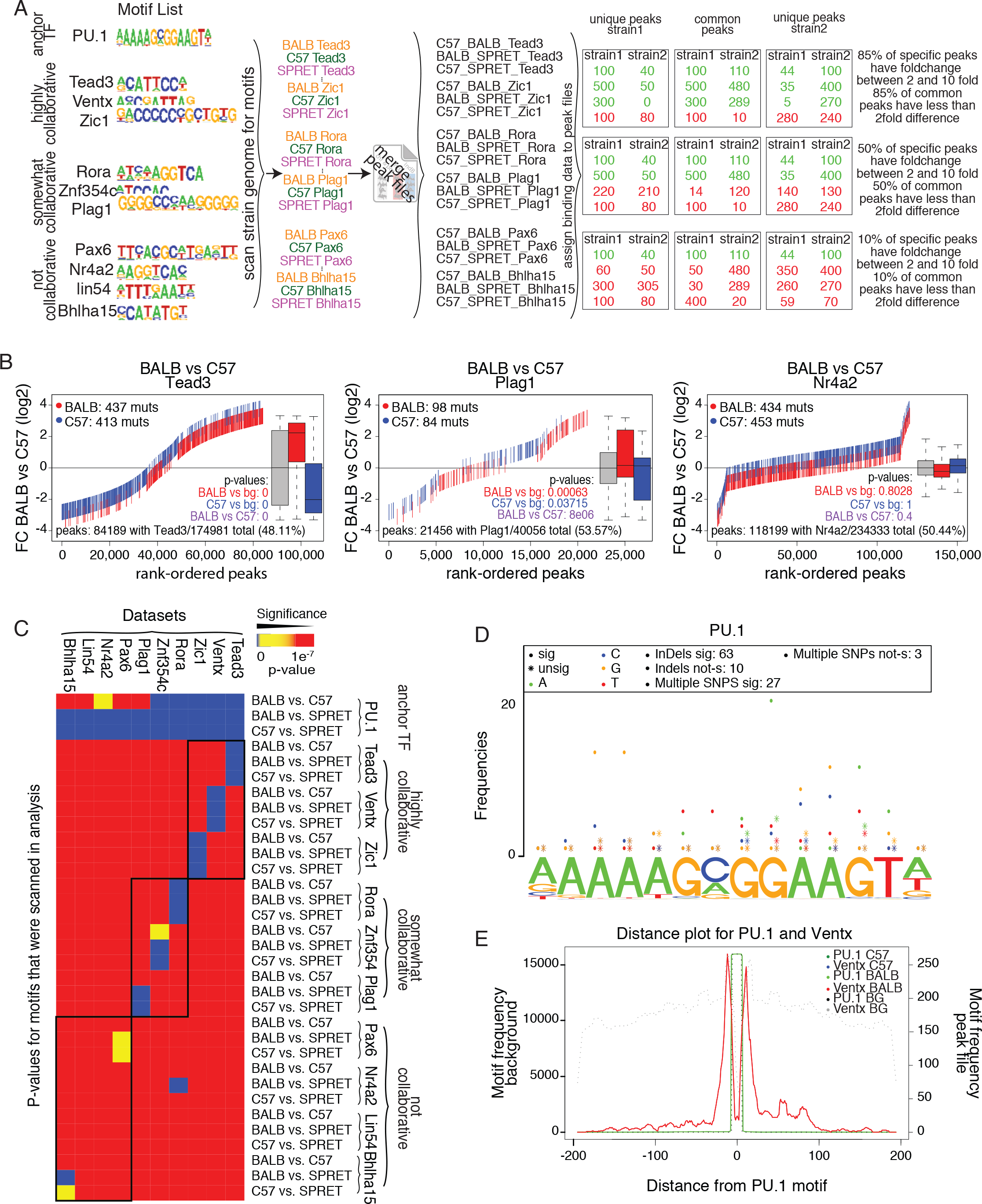
Analysis of a simulated dataset. (**A**) Motifs were defined as important collaborative binding (Tead3, Ventx, Zic1), somewhat collaborative (Rora, Znf354a, Plag1) and not collaborative (Pax6, Nr4a2, lin54, Bhlha15). Peak files were generated for all loci where PU.1 and one of the TF are within 200bp to each other for three mouse strains (C57, BALB, and SPRET) and consecutively merged between two strains. For highly collaborative TF 85% of the strain specific peaks show strain specific binding (somewhat collaborative: 50%, not collaborative: 10%). Fold change was randomly chosen to be between 2 and 10 fold for differently and to be between 1 and 1.5 fold for similarly bound peaks. Read counts were randomly chosen to be between 0 and 500. (**B**) MARGE correctly identifies the association between motif and binding data. Motif mutation distribution plot (Sup. Fig. 2a) for one collaborative motif (Tead3) shows a highly significant association between motif mutation and binding data (medium significance for Plage1 (somewhat collaborative), no significant for Nr4a2 (not collaborative)). (**C**) Summary heatmap for all analysis on the simulated datasets. MARGE showed high significance for the collaborative TF and less or no significance for non-collaborative TF binding motifs. (**D**) Motif mutation position plot for Tead3, showing which positions are mutated and associated with different binding (more information Sup. Fig. 3a). It furthermore shows that in most cases InDels and multiple SNPs cause significant change in binding. (**E**) TF binding motif distribution of PU.1 and Ventx. Motifs for Ventx are closely distributed around the PU.1 binding site (more information Sup. Fig 3b).

### MARGE analysis output

In order to learn more about important position in the motif of the candidate transcription factor, MARGE offers a motif mutation position analysis (Fig. 4d, Sup. Fig. 3a). Fig. 4d shows an example for mutations within the PU.1 motif for the comparison C57 versus BALB on the simulated data set for Ventx. Mutations with significant effects on binding are marked by dots, whereas stars mark mutations with non-significant effects. Each base is colored differently, so it is not only possible to see which positions are mutated (significantly and non-significantly), but also to which other base. In the simulated dataset, even highly conserved residues in the motif can have mutations without an effect on binding (e.g. Fig. 4d, the highly conserved guanine at position 8 has 21 mutations from G‑>A that are significant but also 5 mutations from G‑>A with no effect). In the simulated data this was inherently part of it due to the modeling of biological noise (15% of genotype-specific peaks did show genotype-similar binding). It also should be noted that most differences that could be found were InDels (63 significant versus 10 not significant) or multiple SNPs within one motif (27 significant versus 3 not significant). MARGE also provides a plot that shows the distribution of the Ventx motif around the anchor transcription factor motif PU.1 (Fig. 4e, Sup. Fig. 2b) to see if the motif overlaps the anchor TF motif or if it is only randomly distributed within the peak. This plot allows the user to explore how the motifs of interest are distributed around the center of the peak to get a better understanding of the effect of this motif on the binding of the anchor TF.

### Pairwise analysis of mouse data

To show that the method also works on real data we analyzed data previously published in (21) and (46). We assessed PU.1 (a macrophage LDTF) binding in large peritoneal macrophages (LPM) in three different inbred mouse strains C57BL/6J (C57), NOD/ShiLtJ (NOD), and SPRET/EiJ (SPRET). These strains differ substantially in mutations to each other (Table 1). To show the correctness of the method we generated a list of motifs that were previously discovered (21) to be involved in the establishment of PU.1 binding in macrophage (PU.1, PU.1-IRF, ETS1, SpiB, CEBP, AP-1, Arid3a). Additionally, we chose some transcription factors not expressed in LPMs or with known binding patterns different from PU.1 in macrophages. We chose the motifs of Bcl6 (not expressed in LPM, with a known function in B cells (47)), NeuroD1 (not expressed in LPM, associated with neurons (48) and diabetes (49)), RORgt (not expressed in LPM and mainly associated with thymocytes (50,51)), and Gfi1b (not expressed in LPM and associated mainly with neutrophil differentiation (52)).

MARGE could reliably detect motifs that are significantly associated with PU.1 binding, independent of the number of peaks containing the motif, or the number of mutations in these peaks. For example mutations in CEBP, an important LDTF in macrophages, were detected as significantly associated with PU.1 binding (Fig 5a). The plot showing the positions of mutations within the motif shows enrichment for mutations in the conserved bases T (bases 2 and 3) and A (bases 8 and 9) in comparison to the rest of the bases in the motif (Fig. 5b). Most causal mutations are due to multiple SNPs or InDels, not merely one single SNP. The CEBP motif is distributed closely around the PU.1 motif (where PU.1 is bound) without any motifs overlapping the PU.1 binding site (Fig. 5c). Although the peaks are 200 base pairs with regard to the reference genome, the sequences analyzed can be longer due to long insertions in the different strains resulting in peaks with a size of 300 in this case. Figure 5d shows two examples of how SNPs can influence observed PU.1 binding. In the left panel PU.1 is only bound in SPRET. A SNP in SPRET in comparison to C57 and NOD adds a PU.1 binding motif adjacent to an existent CEBP motif resulting in the observed genotype-specific binding. The right panel shows how loosing a CEBP binding motif in C57 and SPRET close to a PU.1 binding motif existing in all three strains can cause PU.1 binding to be lost. MARGE could not find any significant association between motif existence and binding for the motifs chosen to provide negative controls (Fig. 5e). Although the number of mutations between two genotypes correlates with the significance of the analysis result (due to a bigger sample size), even with a low number of genetic variations MARGE was able to detect almost all significant motifs. To further test MARGE, we applied it to ChIP-seq experiments in four different strains (C57BL/6J (C57), A/J (AJ), CAST/EiJ (CAST), and SPRET/EiJ (SPRET)) for three different factors (CEBPa, FOXA1, and HNF4A) in whole liver from (46). CEBPa is an important TF in hepatocytes (54,55) (which make up about 70% of all cells in the liver (56)) and macrophages. FOXA1 plays important roles for the development and maintenance of the liver, mainly in hepatocytes (15,57) and HNF4A is an important liver TF mainly associated with hepatocytes (reviewed in (58)). Figure 5f shows an example where the TF binding motifs for all three factors were found, but binding could only be observed in AJ, C57, and CAST. Binding in SPRET was lost due to the loss of an adjacent RORA motif. After applying MARGE to the data, all significant motifs were compared to each other and summarized (compare Materials & Methods). In almost all pairwise comparisons for the three different factors the measured factor and the two collaborative factors were found as highly significant (Fig. 5g). Nuclear receptors, which play important roles in the liver (reviewed in (59)), were found as significant in all three comparisons.

**Figure 5:**
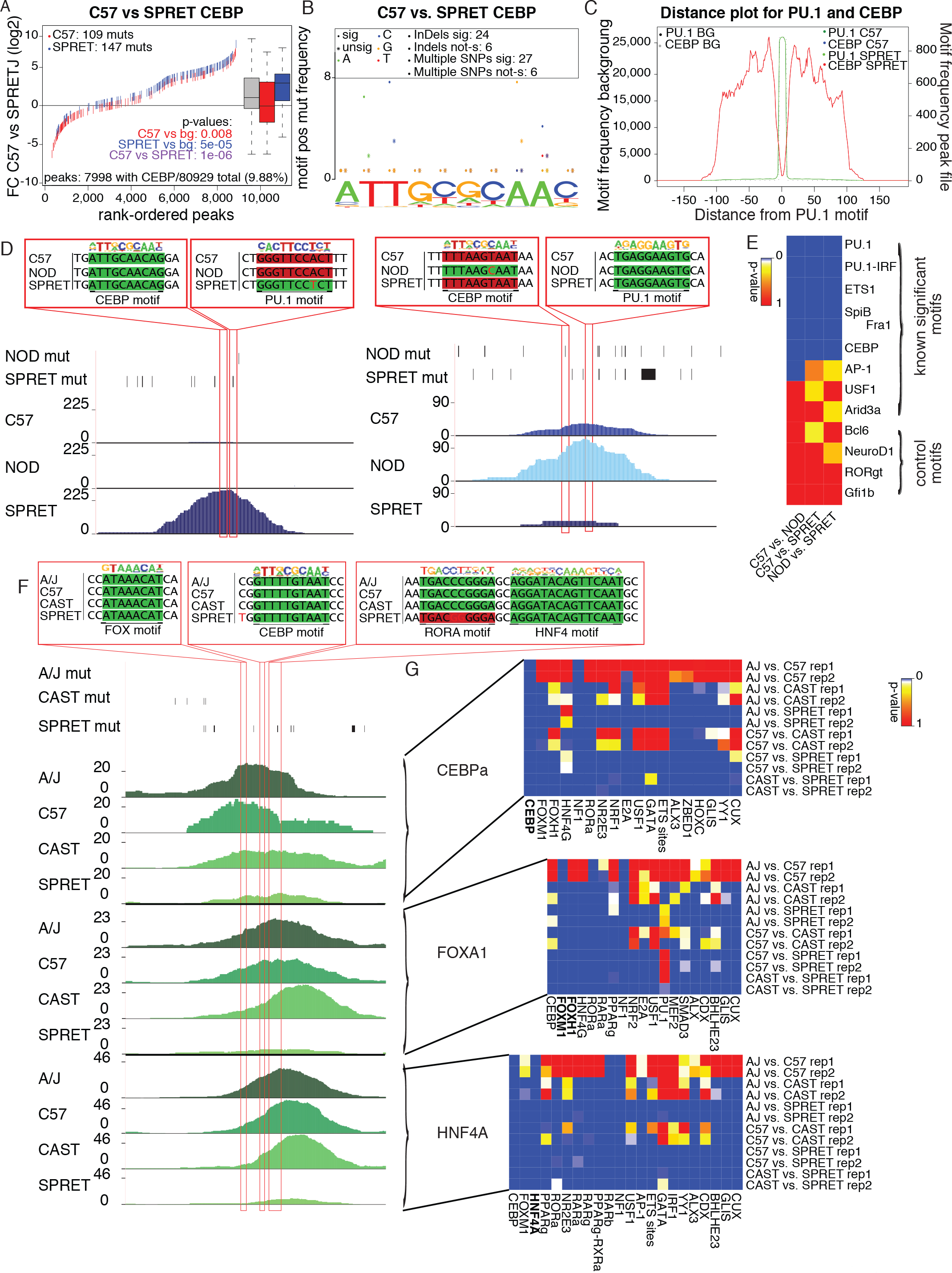
Analysis using MARGE’s pairwise-comparison. (**A**) Motif mutation plot for PU.1 data in LPMs analyzing the impact of mutations in the CEBP binding motif on PU.1 binding. Red ticks show mutations in the CEBP motif in C57 (blue for SPRET). Loss of the CEBP motifs is significantly associated with strain-specific PU.1 binding. (**B**) Motif position mutation plot for CEBP motif showing the position and effect of mutations in the CEBP motif in relation to PU.1 binding. The most conserved positions in the CEBP motif are associated with a loss of PU.1 binding. (**C**) The CEBP motif is distributed closely around the PU.1 motif with a depletion of the CEBP motif at the PU.1 binding site. (**D**) UCSC genome browser shot - Left panel: The gain of a PU.1 motif in SPRET adjacent to a CEBP motif results in PU.1 binding only in SPRET, but not in C57 or NOD. Right panel: The gain of a CEBP motif in NOD in close vicinity to a PU.1 motif results in PU.1 binding only in NOD. (**E**) Summary heat map of multiple testing corrected p-values for TF motifs associated with PU.1 binding. The heat map includes some negative control motifs that are not associated with macrophage biology which were not identified as significant. (**F**) UCSC genome browser shot - The loss of a RORA TF motif in SPRET causes loss of binding of CEBPa, FOXA1, and HNF4A in SPRET, but not in AJ, C57 and CAST. (**G**) Summary heat map of multiple testing corrected p-values of TF binding motifs associated with CEBPa, FOXA1, and HNF4A binding in whole liver. All factors reached significance in every pairwise comparison. Nuclear receptors were significantly associated with binding of the different factors.

### All-versus-all analysis of homozygous mouse data

To show the correctness of the all-versus-all analysis, we reanalyzed the mouse ChIP-seq datasets for CEBPa, FOXA1, and HNF4A from whole liver (Fig. 6a). Almost all motifs that were found significant in at least one pairwise comparison were detected as significant in the all-versus-all comparisons (compare Fig. 5g and Fig. 6a). Applying the motif score or the motif existence in the LMM produced almost the same results, with some motifs differing. The motif existence approach should be used with caution since adjusting the threshold that defines a sequence as motif can have large impacts on the results. Therefore, the all-versus-all comparison is able to confirm motifs significantly associated with binding of CEBP, FOXA1, or HNF4A in whole mouse liver previously identified by MARGE’s pairwise comparisons. To make sure that the all-versus-all comparison is sensitive, we shuffled the strain order and repeated the analysis (Fig. 6b). To assess how much the results are influenced when very similar strains are shuffled, AJ and C57 were switched, but CAST and SPRET were kept at the same position. The further assess robustness of the results, the more diverse strains were shuffled with the more similar strains. Furthermore, we used completely different mouse genomes (NOD, DBA, PWK, and WSB). The color bar in Figure 6b shows the number of differences between the strains. When two very similar strains were changed (AJ with C57) the results are almost the same and the data sets are clustered together. However, as soon as more different strains are switched, the results changed dramatically. Motifs that are significant in all comparisons (e.g. NF1) should be counted as false positive results. This analysis shows that changing very similar data sets with each other does not affect the results, probably because most of the informative loci are found between these two strains and the two more diverse strains.

**Figure 6:**
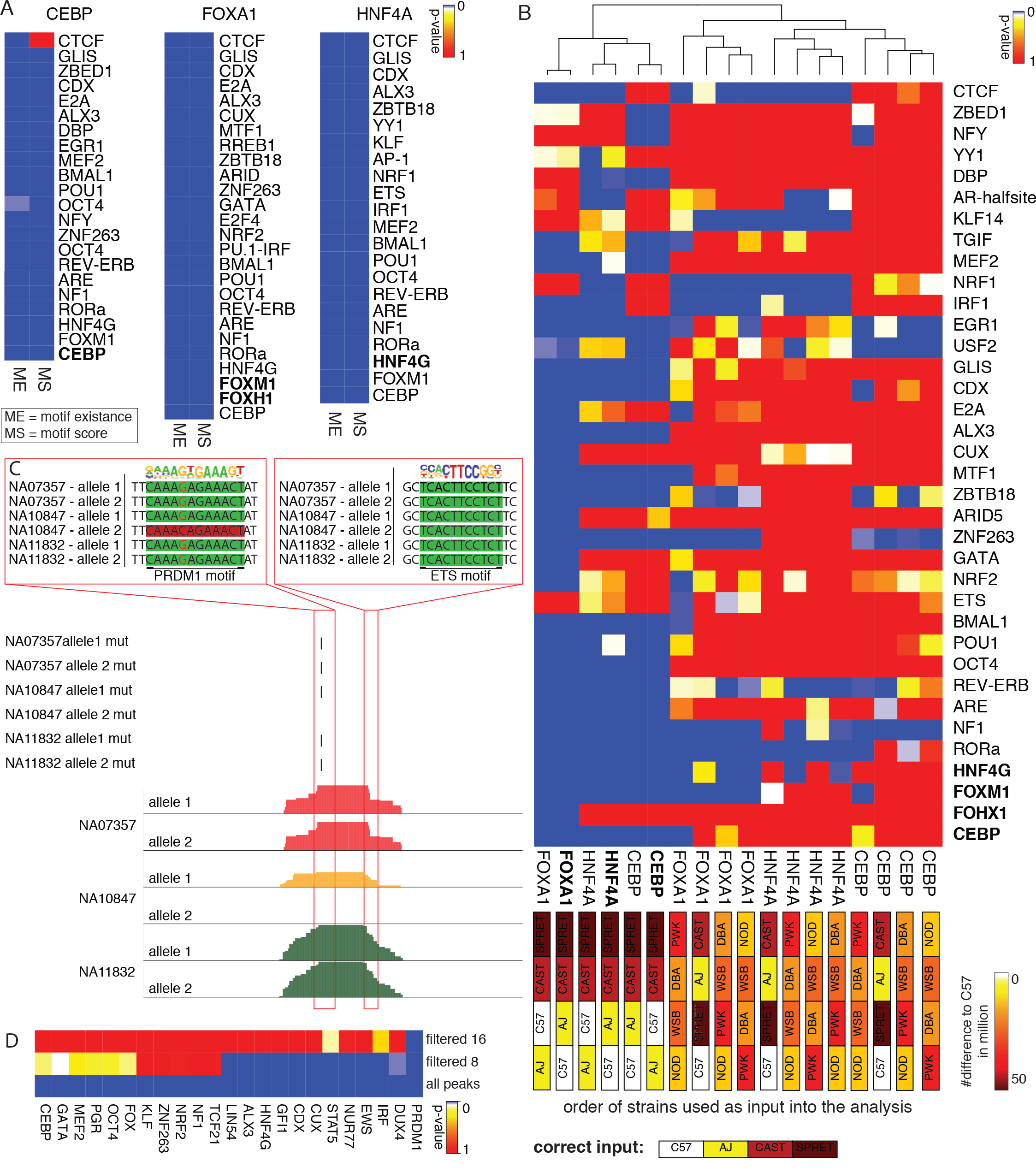
Results of all-versus-all analysis. (**A**) Summary heatmap of multiple testing corrected p-values of all-versus-all analysis of CEBP, FOXA1, and HNF4A ChIP-seq data sets from whole liver in AJ, C57, CAST, and SPRET. The analysis confirms the results from the pairwise analysis performed in Fig. 5g. The same motifs are highly significance with slight variations independent of considering motif score (MS) or motif existence (ME). (**B**) Summary heatmap of multiple testing corrected p-value of the all-versus-all analysis for CEBP, FOXA1, and HNF4A ChIP-seq data sets with the original order of the strains and shuffled order of the strains to assess sensitivity of MARGE. The color of the boxes correlates to the number of mutations (from 0 - white to 50 million - brown). When very similar strains are switched (AJ and C57) the MARGE results are clustered together. As soon as more diverse strains are switched or different strains are used, the results cluster as outliers to the original data and almost all motifs lose significance. (**C**) UCSC genome browser shot visualizing three human PU.1 datasets. The allele-specific loss of a PRDM1 motif close to an ETS motif causes allele-specific loss of PU.1 binding. (**D**) Summary heat map of multiple testing corrected p-values of transcription factor motifs significantly associated with PU.1 binding in human lymphoblastoid cell. Many TF motifs found to be significantly associated with PU.1 binding are either known to play important roles in B cell development and maintenance or cancer. By increasing the stringency of peaks included in the analysis (and decreasing the number of observations) the number of significant motifs decreases. Only PRDM1 is found as significant when using a filter of 16 reads.

### All-versus-all analysis of heterozygous human data

To show that MARGE is also able to analyze data from several human individuals with a low number of mutations, 34 PU.1 ChIP-seq datasets from Waszak at el (25) were analyzed with MARGE (listed in Sup. Table 1). The VCF files were downloaded from the 1000 Genomes Project (30) and the individual MARGE files and genomes were generated. A bowtie2 (27) index was created for each genome (two indices per genotype - one for the complete genome containing mutations on allele 1 and one for mutations on allele 2) and the ChIP-seq reads were mapped against both indices of the corresponding genotype. Only data sets with an overall mappability of 80% were considered in the downstream analysis (22 individuals) Peaks were called on all perfectly aligned reads and all peaks were merged and annotated allele-specific (320,146 peaks). To see how noise influences the MARGE results, MARGE was applied to an unfiltered peak file, as well as a peak file only containing reliable peaks with at least 8 reads in at least on individual (16, respectively). The dataset used in this analysis was based on lymphoblastoid cell lines, human B cell lines infected with an Eppstein-Barr virus to immortalize them.

Because the dataset is based on B cells it is not expected that any macrophage specific LDTFs are significant, instead B cell specific LDTFs (like PRDM1 also known as BLIMP-1, E2A etc.) would be expected to show a significant association with PU.1 binding (60). Figure 6c shows a UCSC genome browser session for one locus in three different individuals where one SNP that causes a loss of a PRDM1 motif close to an ETS factor motif is associated with loss of binding of PU.1. Applying the mutation approach systematically to all loci in all individuals and then summarizing the motifs, MARGE identified the B cell LDTF PRDM1 as highly significant, as well as a motif belonging to the IRF family of transcription factors known to play a role in B cells (Fig 6d) and an ETS motif, important for PU.1 binding. DUX4 has been previously associated with acute lymphoblastic leukemia (ALL) (61) which is coherent with the cancer-like cell type used in this experiment. MARGE was able to identify many other important transcription factors for B cells including NUR77 and a KLF binding motif (associated with B cell development (62,63)). The more stringent the filtering, the less significant motifs could be found. Filtering by 8 reads, about half of the significant motifs could be found. But filtering by 16 reads only found PRDM1 as significant. This highlights the importance of a good quality data set, because a lot of difference is found in lower bound peaks rather than the top peaks. Overall, MARGE was able to find significant motifs associated with PU. 1 binding in human lymphoblastoid cell lines taking advantage of allele-specific binding in many individuals.

## Discussion

We developed a powerful tool to efficiently analyze ChIP-seq and other NGS data to understand the impact of transcription factor motifs on collaborative binding of transcription factors. MARGE is the first publicly available suite of software tools to integrate natural genetic variation (including InDels) and NGS binding data and provides complementary algorithms to analyze data from different genetic backgrounds in a pairwise manner as well as by utilizing a linear mixed model. It further provides many useful tools to directly look at genetic differences between different genetic backgrounds. By simulating a dataset and also applying MARGE to real world data, we could show that the algorithm works correctly in identifying motifs significantly associated with the binding of a measured transcription factor. Here, we applied MARGE to ChIP-seq data, which requires a well-working antibody for the reference transcription factor. However, MARGE can also be applied to ATAC-seq data or DNase I hypersensitivity data, which does not require any previous knowledge. In this case, rather than collaborative binding partners for a reference transcription factor, analysis of open chromatin would be expected to recover the dominant collaborative factors needed to establish open chromatin regions. Therefore, MARGE can potentially be applied to identify key regulatory factors in any cell type as long as parallel datasets from genetically diverse strains or individuals are available.

The algorithm assumes that the binding of the measured factor is only affected by local mutations in transcription factor binding motifs. As a result, sequence changes that influence binding on a global or long-distance scale in *trans* will not be detected and introduce noise to this. Furthermore, MARGE only analyzes one motif at a time. More complex relationships between transcription factors (e.g. the requirement for binding of three factors simultaneously) are not considered in the analysis. As in every analysis based on statistical tests, the power of discovery is dependent on the number of observations. A greater number of genetic variations between two individuals provides a better analysis result and will detect more significant motifs. For comparisons with low numbers of genetic variations MARGE offers a linear mixed model to increase the power of detection by merging all genetic variation between all individuals. This, however, requires substantially more experiments. Furthermore, the software is dependent on a list of position-weight matrices for the detection of TF binding sites. It is known that TF can bind to very weak motifs that cannot be detected by a motif-scanning algorithm but play important roles in regulating gene expression (64). However, MARGE is dependent on finding motifs based on scanning the DNA for the consensus sequence provided by the PWM. This limits the sensitivity of MARGE. Improvements in our understanding how to detect motifs in sequence will therefore improve the power of MARGE. Similar to de-novo motif finding, also MARGE only detects TF motifs. There are sometimes many similar transcription factors capable of binding the same consensus motif, which MARGE cannot discriminate. As more TFs and their motifs are characterized, these types of analysis will surely improve.

Genome-wide association studies (GWAS) evaluating common sequence variants associated with diverse phenotypes consistently demonstrate that the majority of variants reside in noncoding regions of the genome (20,65,66). These findings suggest that such variants impose risk by altering promoter and enhancer elements that regulate gene expression. Interpretation of such variants is currently limited because the genomic location of the regulatory elements at which they could potentially exert their effects varies according to cell type. By identifying important motif mutations, MARGE can provide a new and unique way to analyze transcription factor binding and detect the major collaborative factors involved in the establishment of cell-specific enhancer landscapes. With the advances in sequencing technology and availability of human samples, MARGE can facilitate the analysis of datasets that provide insights into the relationship between non-coding genetic variation and gene expression in humans.

## Availability

The MARGE source code and installation package are freely available on GitHub: https://github.com/vlink/marge/blob/master/MARGE_v1.0.tar.gz.

The mouse LPM dataset from (21) was downloaded from the GEO database under accession number GSE62826. The data is available at http://genome.ucsc.edu/cgi-bin/hgT_racks?hgS_doOtherUser=submit&hgS_otherUserName=vlink&hgS_otherUserSessionName=MARGE_LPM_data. The mouse liver data set from (46) was downloaded from ArrayExpress Archive (http://www.ebi.ac.uk/arrayexpress/) under accession number E-MTAB-1414. The data is available at http://genome.ucsc.edu/cgi-bin/hgT_racks?hgS_doOtherUser=submit&hgS_otherUserName=vlink&hgS_otherUserSessionName=MARGE_Liver_data. The human data set from (25) was downloaded from the ArrayExpress Archive under accession number E-MTAB-3657. The data is accessible at http://genome.ucsc.edu/cgibin/hgTracks?hgS_doOtherUser=submit&hgS_otherUserName=vlink&hgS_otherUserSessionName=MARGE_Liver_data..

MARGE is implemented in Perl and R (44). It has been tested on several UNIX systems, including CentOS and Debian with Perl version 5.20 and higher and R version 3.3 and higher. We provide a script that installs MARGE and allows download of pre-processed mutation data from the mouse genome project (29) and the genomes from the 1000 Genome Project used in this manuscript. MARGE requires the Perl core modules POSIX, Getopt::Long, Storable and threads, as well as the modules Set::IntervalTree (67), and Statistics-Basic (68). It further requires the R packages SeqLogo (69), gridBase (70), lme4 (71), and gplots (72). It also requires an installed version of gzip. For the motif mutation analysis MARGE requires HOMER (1) (http://homer.ucsd.edu/homer/) to be installed and executable. Without a working installation of HOMER, MARGE’s functionality is limited to only visualization and annotation of the data.

## Acknowledgements

We thank Leslie van Ael for support with figures. We further thank Ty Troutman, Jenhan Tao, and Inge Holtman for running MARGE and their feedback. We thank Chris Benner, Ty Troutman, Dylan Skola, Jenhan Tao, and Zhengyu Ouyang for critically reading the manuscript.

---

**Supplemental Figure 1:**
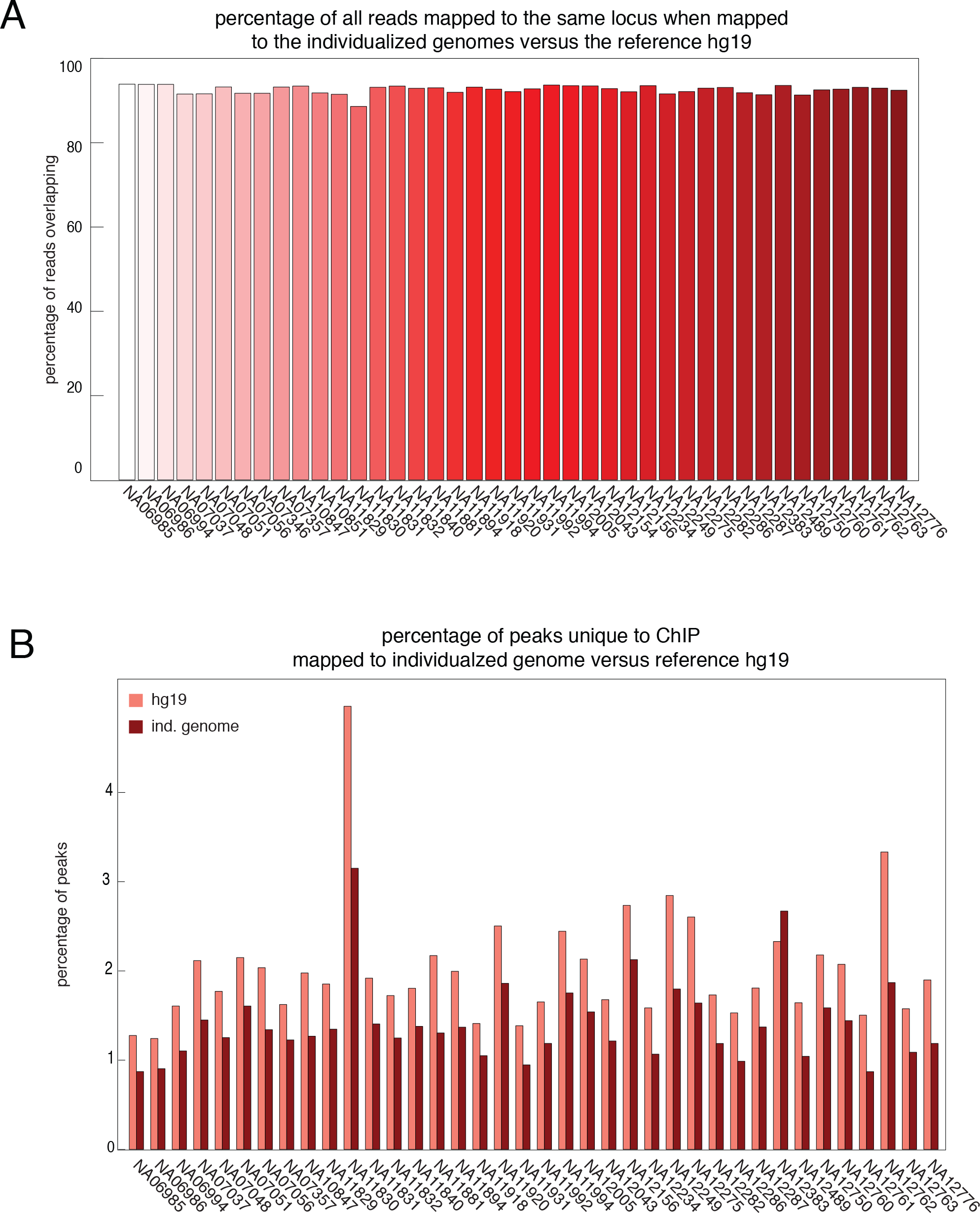
Effect of mapping ChIP-seq data for human lymphoblastoid cell lines. (**A**) Percentage of reads mapped to the same locus after mapping to individualized genotype and reference. Only about 90% of reads mapped to the same locus when comparing mapping results for individualized genomes versus the reference. (**B**) Percentage of peaks unique to either the individualized genome of the reference genome hg19. Up to 4% of the peaks were called uniquely in either the PU.1 ChIP-Seq dataset mapped to hg19 or the individualized genome.

**Supplemental Figure 2:**
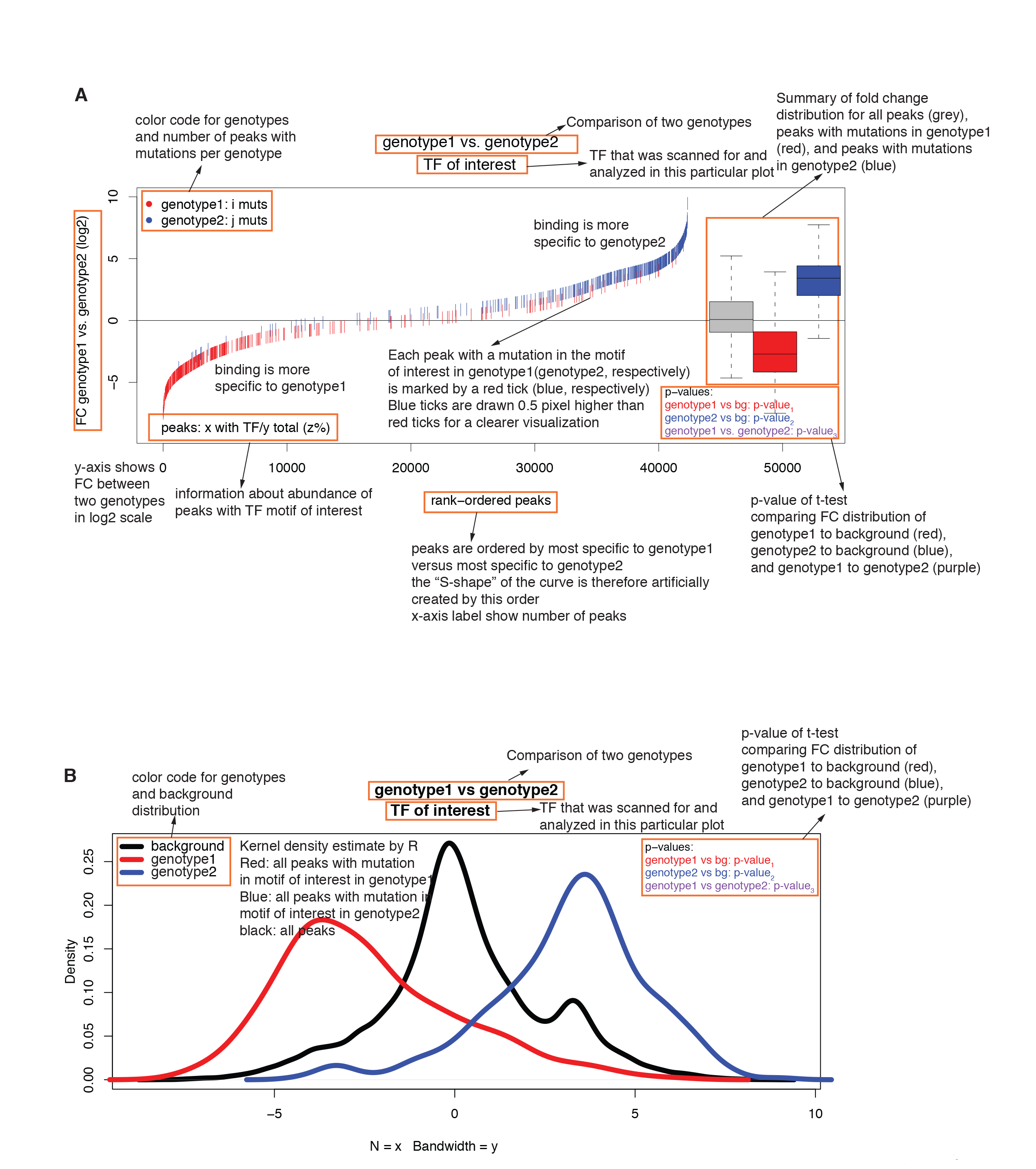
Detailed description of the output plots MARGE generates for the motif mutation analysis. (**A**) For each TF of interest MARGE generates this plot. The peaks are rank-ordered by most genotype-specific bound in genotype1 to most genotype-specific bound in genotype2. Genotype1 is color-coded red, whereas genotype2 is color-coded blue. The left upper corner shows how many mutations could be found in the motifs for genotype1 and genotype2. A red tick marks each peak without a TF binding motif of the TF of interest, if the motif is missing in genotype1 and a blue tick marks the motif is missing in genotype2. All data on the left bottom of the plot show peaks where the binding is very specific to genotype1 The right upper corner shows peaks where binding is very specific to genotype2. The box plot on the right summarizes the fold change distribution. Grey shows the fold change distribution for all peaks having the TF binding motif of interest. Red shows the fold change distribution for all peaks missing the TF binding motif of interest in genotype1, whereas blue shows the distribution for all peaks missing the TF binding motif in genotype2. A Student’s t-test is performed comparing these distributions. The p-value is shown below the box plots. Red shows the comparison of the background (grey box) versus the distribution for genotype1 (red box), blue shows the comparison of the background (grey box) versus the distribution for genotype2 (blue box) and purple shows the comparison between genotype1 (red) and genotype2 (blue). (**B**) Kernel density plot for data shown in the motif mutation distribution plot. A Gaussian kernel is applied to the fold change distributions for all peaks with the motif of interest (black - background), all peaks with missing motifs in genotype1 (red) and all peaks with missing motifs in genotype2 (blue) and plotted. The p-values are the corresponding p-values from the t-test explained above.

**Supplemental Figure 3:**
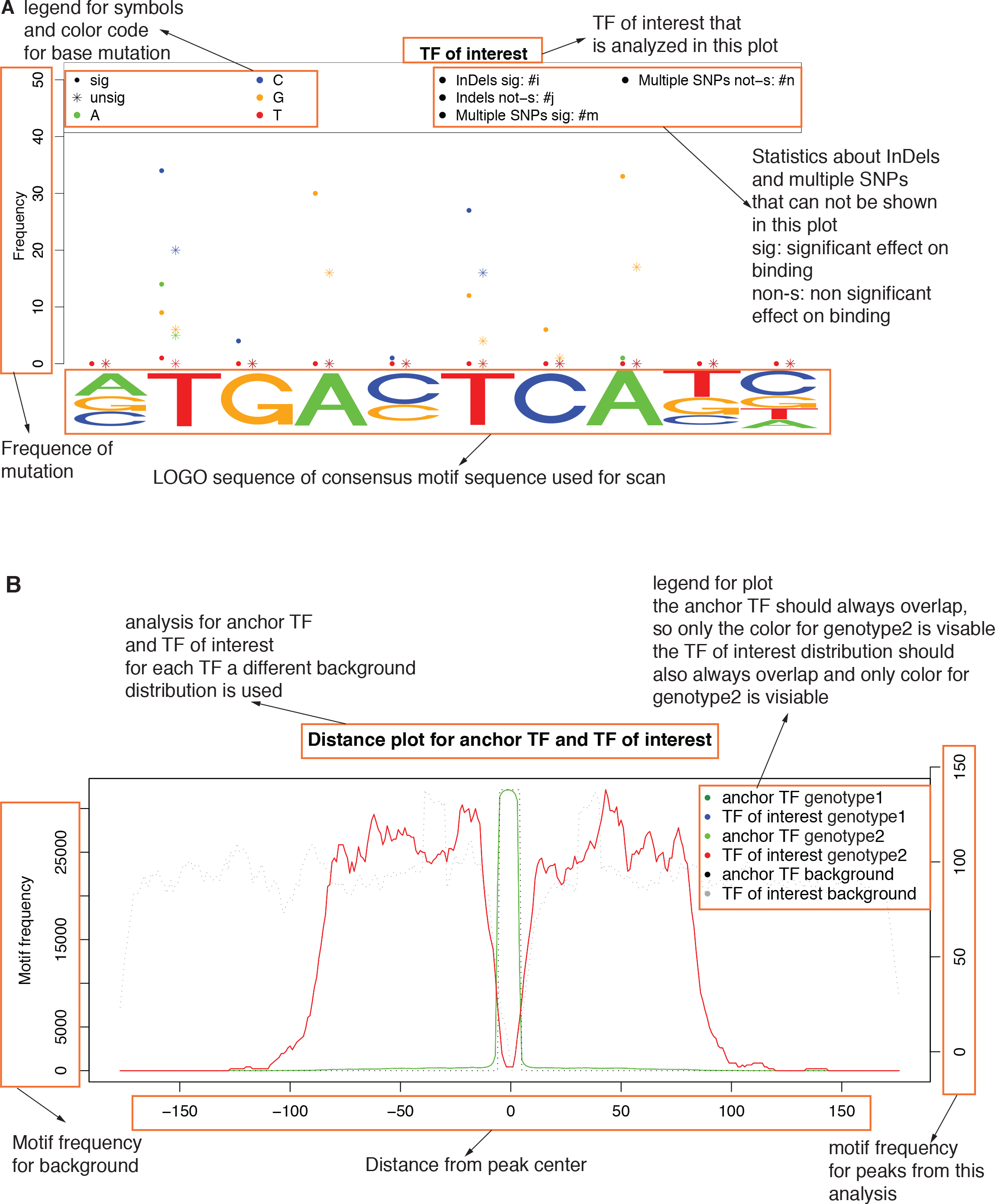
Detailed description of additional output plots provided by MARGE. (**A**) Motif position mutation plot shows the consensus logo of the position-weighted matrix (PWM) from the motif. Mutations that affect the TF binding between both genotypes are marked by dots, whereas a star marks mutations with no effect. All mutations resulting in an adenosine are color-coded green (cysteine are color-coded blue, guanine are yellow and thymine are red). The number of InDels and multiple SNPs are reported separately for significant changes in binding and no changes. The y-axis shows the frequency for the mutations. (**B**) Motif distance distribution plot. All peaks are centered on their anchor TF binding motif and the distribution of the TF of interest is plotted around the TF. Further, a genome-wide background is plotted. The right y-axis shows the motif frequency in the genotypes, the left y-axis shows the frequency in the background. The distribution of the anchor TF is plotted for the background and the genotypes in the order of background (grey), genotype1 (dark green), and genotype2 (light green). They should overlap, so only the distribution of genotype2 should be visible. For the motif of interest the genome wide background is plotted in grey, whereas the distribution for genotype1 is blue and for genotype2 is red. The distribution should overlap and only the distribution of genotype2 should be visible. The x-axis shows the distance of the motif to the center.

**Supplemental Table 1:** Summary of peak numbers before and after shifting for all 34 individuals from a human PU1 ChIP-Seq dataset. Only up to 11 peaks are lost after shifting, which is less than 0.1% of all peaks.

**Supplemental Table 2:** Summary of peak numbers before and after shifting for mouse data. The number of genetic variations is up to 40 million between two strains, and less than 0.1% of the peaks are lost after shifting.

